# Inactivity and Ca^2+^ signaling regulate synaptic compensation in motoneurons following hibernation in American bullfrogs

**DOI:** 10.1101/2022.01.26.477843

**Authors:** Tanya Zubov, Lara do Amaral-Silva, Joseph M. Santin

## Abstract

Neural networks tune synaptic and cellular properties to produce stable activity. One form of homeostatic regulation involves scaling the strength of synapses up or down in a global and multiplicative manner to oppose activity disturbances. In American bullfrogs, excitatory synapses scale up to regulate breathing motor function after inactivity in hibernation, connecting homeostatic compensation to motor behavior. In traditional models of homeostatic synaptic plasticity, inactivity is thought to increase synaptic strength *via* mechanisms that involve reduced Ca^2+^ influx through voltagegated channels. Therefore, we tested whether pharmacological inactivity and inhibition of voltage-gated Ca^2+^ channels are sufficient to drive synaptic compensation in this system. For this, we chronically exposed *ex vivo* brainstem preparations containing the intact respiratory network to tetrodotoxin (TTX) to stop activity and nimodipine to block L-type Ca^2+^ channels. We show that hibernation and TTX similarly increased motoneuron synaptic strength and that hibernation occluded the response to TTX. In contrast, block of L-type Ca^2+^ channels did not alter mean synaptic strength but disrupted the apparent multiplicative scaling of synaptic compensation typically observed in response to hibernation. Thus, inactivity drives up synaptic strength through mechanisms that do not rely on reduced L-type channel function, while Ca^2+^ signaling associated with the hibernation environment independently regulates the balance of synaptic weights. Altogether, these results point to multiple feedback signals for shaping synaptic compensation that gives rise to proper network function during environmental challenges *in vivo.*

## Introduction

Neural circuits must produce correct output patterns to generate reliable behaviors. However, animals face constantly changing internal and external environments, posing a threat to stable neuronal function^1^. To maintain stability, the nervous system is thought to use a wide range of homeostatic mechanisms that sense activity perturbations which then modify cellular and synaptic function to regulate neuronal output. Although diverse homeostatic mechanisms exist^2,3^ the most well-studied involves compensatory changes in excitatory synaptic strength through a mechanism termed synaptic scaling. Synaptic scaling is interpreted to involve the up or downregulation of all a neuron’s synaptic inputs by the same relative amount to oppose activity disturbances^4^ Mounting evidence implicates synaptic scaling in association with diverse physiological conditions such as development^5,6^ sleep^7^, and learning^8^ in mammals, while inappropriate function of homeostatic mechanisms may lead to various neurological disorders^9–11^. Although a large body of work supports cell-wide multiplicative synaptic compensation^4,5,8,12–15^ how these processes are regulated and integrated *in vivo* to generate adaptive behaviors is less clear.

Animals that hibernate must enter and exit states of long-term dormancy, making excellent models to connect activity-dependent neural compensation to behavior Motor circuits that produce breathing in American bullfrogs, *Lithobates catesbeianus,* are especially intriguing in this regard. The respiratory network typically generates stable rhythmic motor activity throughout life. However, during under water hibernation, gas exchange demands are met through cutaneous respiration, and the motor circuits that produce lung breathing stop entirely^17–19^ After months of inactivity, frogs emerge in the spring and quickly display normal respiratory motor performance^17,20^ We previously found that motoneurons scale up excitatory synapses in response to hibernation, which acts to maintain respiratory motor outflow immediately after the animal restarts breathing^15^ Therefore, this system allows the study of synaptic compensation in relation to motor behavior and points to conserved mechanisms across mammals and amphibians in the natural environment.

In mammalian systems, synaptic compensation is thought to be induced by variables related to inactivity, for example, decreases in firing rate or postsynaptic receptor activation^21–23^ From there, decreases in intracellular calcium represents a message that is proportional to activity, with the capacity to influence Ca^2+^-dependent signaling that drives compensation^24^ Reduced Ca^2+^ influx through L-type calcium channels in particular plays a critical role in eliciting synaptic compensation in diverse neuronal types^22,25,26^ Given that hibernation causes a large and chronic reduction in neuronal output, inactivity and subsequent reductions in Ca^2+^ influx may trigger synaptic compensation that serves to regulate motor performance^15,16^ However, aquatic hibernation involves diverse physiological changes to the animal^27^, raising questions about whether inactivity or other aspects of the organismal environment, such as temperature, drive synaptic compensation. To begin to approach this question, we tested the hypothesis that inactivity and reduced voltage-gated Ca^2+^ channel function are sufficient to drive up synaptic strength in frog respiratory motoneurons. The amphibian respiratory network is well-suited to address this question, as “fictive breathing” motor patterns persist in dissected brainstem preparations for >24 hours *ex vivo* This allowed us to perform chronic pharmacological manipulations to the native network without the need for cell culture conditions. To this end, we isolated the respiratory network *ex vivo,* silenced activity with tetrodotoxin (TTX) or blocked L-type Ca^2+^ channels with nimodipine. We then compared synaptic responses to these manipulations with that seen during hibernation and then tested whether hibernation occludes the synaptic response induced by these drugs. This allowed us to infer whether network inactivity and Ca^2+^ channel blockade overlap with pathways associated with hibernation, a similar “sufficiency/occlusion” approach used to relate long-term potentiation of synaptic strength to learning at hippocampal synapses^29^.

## Results

Amphibians inflate the lungs with positive pressure, whereby muscles of the buccal floor and glottis form a pump-valve system for lung ventilation. Laryngeal motoneurons that innervate the glottal dilator, the value muscle, (Fig. 1A) undergo compensatory upregulation of AMPA-glutamate receptor currents in response to ~2 months of aquatic hibernation^15^ To simulate the overwintering environment in this study, frogs were cooled from 20°C to 2°C over one week and then held at 2°C for either two or four weeks before assessing synaptic strength in brain slices. Consistent with our previous report with inactivity at longer time scales, synaptic strength of laryngeal motoneurons, as assessed by the amplitude of spontaneous excitatory postsynaptic currents (sEPSC), increased in frogs exposed to the simulated hibernation environment at both two and four weeks (Fig. 1B-C) with no change in sEPSC frequency (Fig. 1D). These results confirm postsynaptic compensation in response to hibernation and demonstrate that upregulation of excitatory synaptic strength plateaus at a new steady-state value relatively early into aquatic submergence. All subsequent experiments were performed in animals that experienced 4 weeks in the hibernation environment.

**Figure 1.**
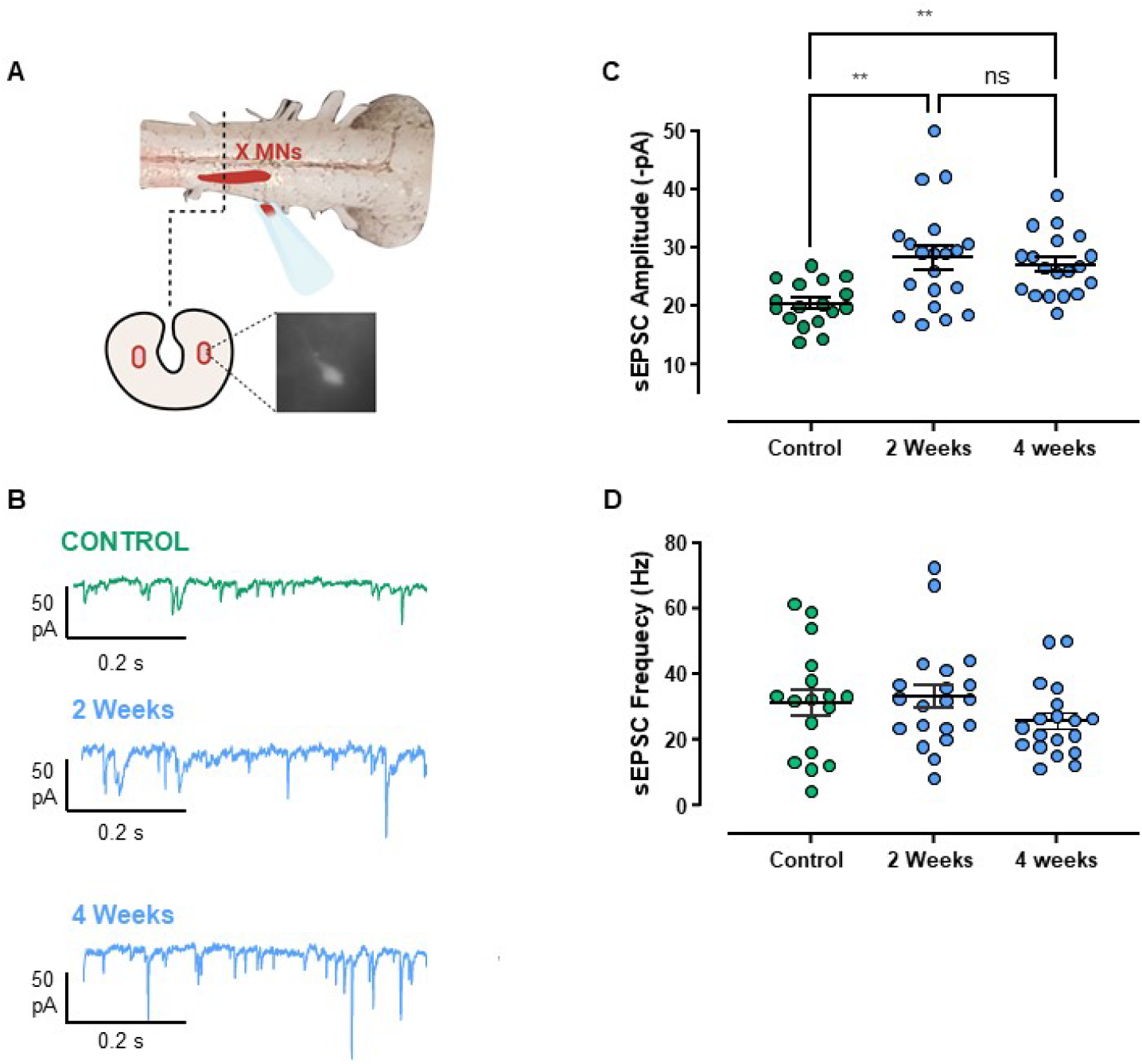
sEPSC amplitude, but not frequency, increases by two weeks of cold-acclimation. (A) Schematic of the experimental process to label respiratory motoneurons. The 4^th^ root of the vagus nerve was backfilled with fluorescent dextran, and neurons were identified in slices through fluorescence imaging microscopy. Animals were either warm-acclimated (control; 22°C; n=17 from 6 animals), or cold-acclimated for two (n=20 from 4 animals) or four weeks (n=19 from 4 animals). (B) Example voltage clamp recordings of sEPSCs in laryngeal motoneurons from controls and frogs that hibernated for 2 and 4 weeks. (C) sEPSC amplitude was increased at 2 weeks and remained elevated at 4 weeks (One-way ANOVA; p=0.0013), but sEPSC frequency was not affected (D, p=0.287, Kruskal-Wallis test). Dots represent data points for individual neurons, and average values are indicated by the horizontal line. Error bars are S.E.M. **p<0.01. Error bars are S.EM.

Variables associated with inactivity *(i.e.,* loss of postsynaptic firing or synaptic transmission) can drive homeostatic increases in synaptic strength^21,22^ Our rationale was that if synaptic compensation during winter is responding to inactivity, pharmacological inactivity may produce a similar degree of synaptic compensation as hibernation, thus demonstrating inactivity as a sufficient signal to generate the hibernation response. To test this, we silenced intact respiratory networks in the brainstem-spinal cord preparation with 100 nM TTX for 16 hours. Fig. 2A shows extracellular recordings of respiratory-related motor output from the vagus nerve, highlighting that activity is present in time control and post-hibernation preparations throughout the 16 hr period but eliminated by TTX. While the time course of hibernation (one month) and shorter-term TTX (16 hours) are different, synaptic scaling has been observed to manifest after 4 hours of inactivity *in vitro* We compared average miniature excitatory postsynaptic current amplitude in slices (mESPCs) following network inactivity induced by TTX to Time Controls (control preparations with rhythmic activity present for 16 hrs at 22°C) and Post-Hibernation preparations (preparations recovered following hibernation for 16 hrs at 22°C; Fig 2A). TTX and hibernation groups exhibited similar mean increases in cell-averaged mEPSC amplitude relative to Time Control preparations. These mean increases were not different from each other (Fig. 2B-C). Therefore, TTX produces a similar average amount of synaptic compensation as hibernation, despite dramatically different durations and modes of inactivity.

**Figure 2.**
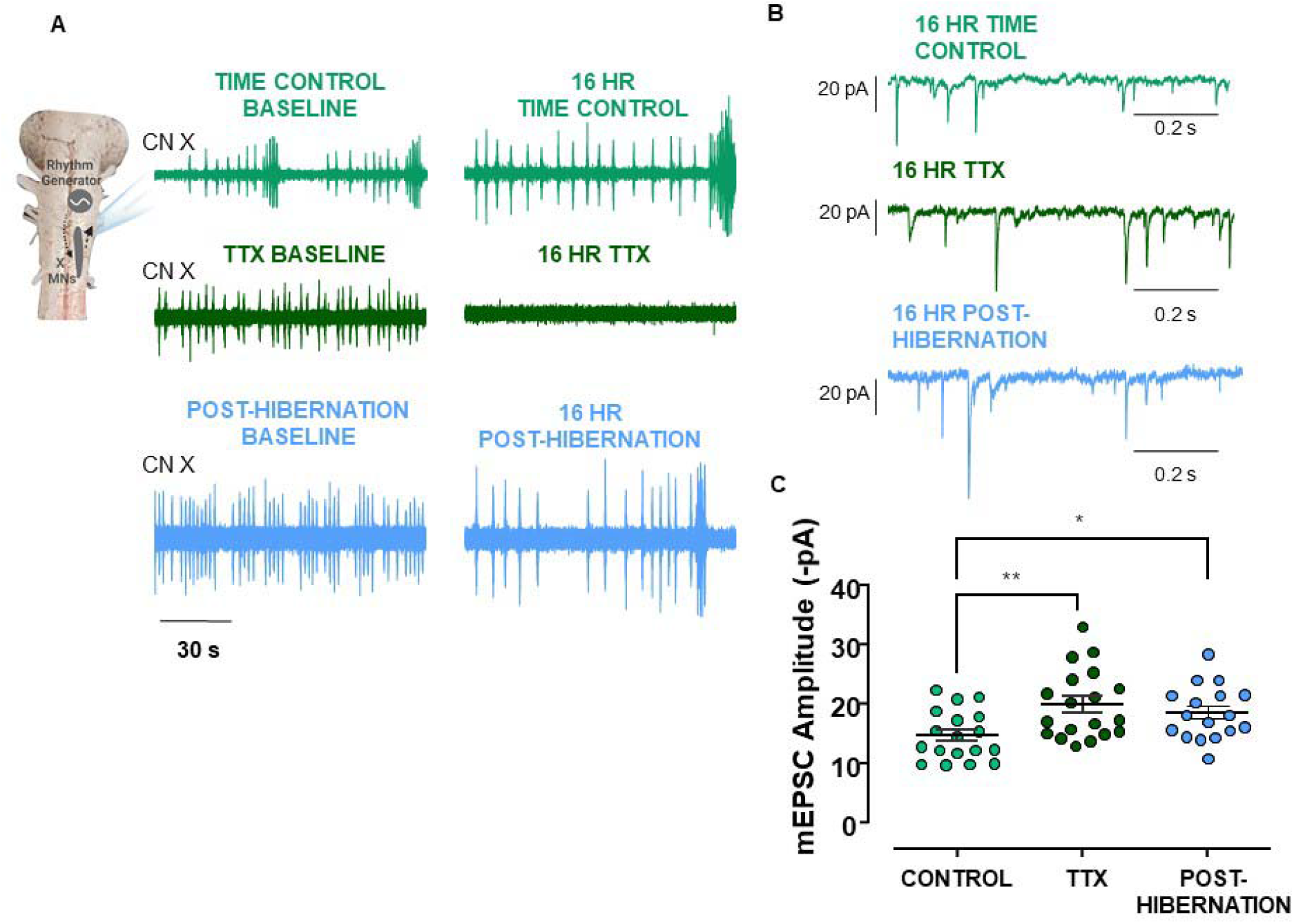
Hibernation and TTX induce increases in mEPSC amplitude to the same extent. (A) Example respiratory-related vagal motor output (CNX) driven by the respiratory rhythm generator, highlighting activity for the entire incubation period. TTX silenced activity in all preparations (B) Example mEPSC traces recorded in whole-cell voltage clamp in laryngeal motoneurons in time control, TTX, and following hibernation. (C) Mean data showing enhanced mEPSC amplitude following both hibernation and TTX exposure relative to time control (One-way ANOVA, Holm-Sidak Multiple Comparisons Test, control vs. TTX, p=0.004, control vs. post-hibernation, p=0.0139, TTX vs. post-hibernation, p=0.4405). Circles represent cell-averaged mEPSCs for each neuron. Error bars are S.E.M. Horizontal line represents the mean. Control (n=18 from 5 animals), TTX (n=19 from 5 animals), post-hibernation (n=17 from 4 animals). *p<0.05, **p<0.01.

The previous experiment demonstrated that inactivity *via* TTX is sufficient to drive synaptic compensation in this system. Next, we determined if hibernation occludes the synaptic response to TTX. If hibernation prevents further increases by TTX, this may reflect overlapping signaling processes. However, if TTX and hibernation produce additive responses, this may suggest divergent signaling pathways for compensation. When we exposed post-hibernation preparations to TTX for 16 hr, we did not observe further increases in synaptic strength by TTX compared to hibernation alone (Fig. 3A-B). Membrane properties, mEPSC frequency, and mEPSC decay time constant were not different across control, post-hibernation, TTX, and post-hibernation+TTX groups (Fig. 4). Overall, these data suggest that hibernation occludes the synaptic compensation response to TTX.

**Figure 3.**
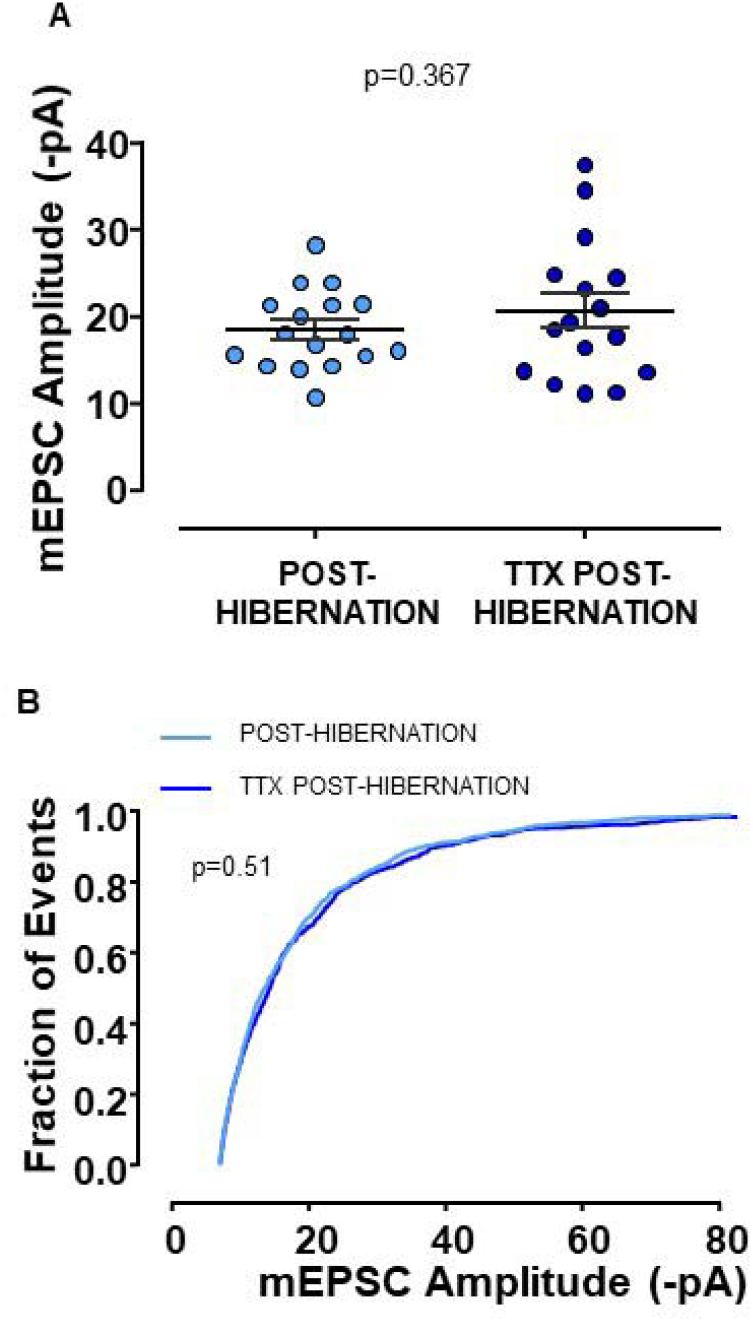
Hibernation occludes the mEPSC amplitude increase by TTX. (A) shows the mean mEPSC amplitude after hibernation and hibernation in the presence of 16 hr TTX (unpaired t test). Post-hibernation, n=17 from 4 animals and TTX Post-hibernation n=16 from 4 animals. (B) shows the cumulative distribution of post-hibernation and TTX post-hibernation mEPSC amplitudes (p value from Kolmogorov-Smirnov Test). Circles represent data points for individual neurons, and average values are indicated by the horizontal line. Error bars are S.E.M.

**Figure 4.**
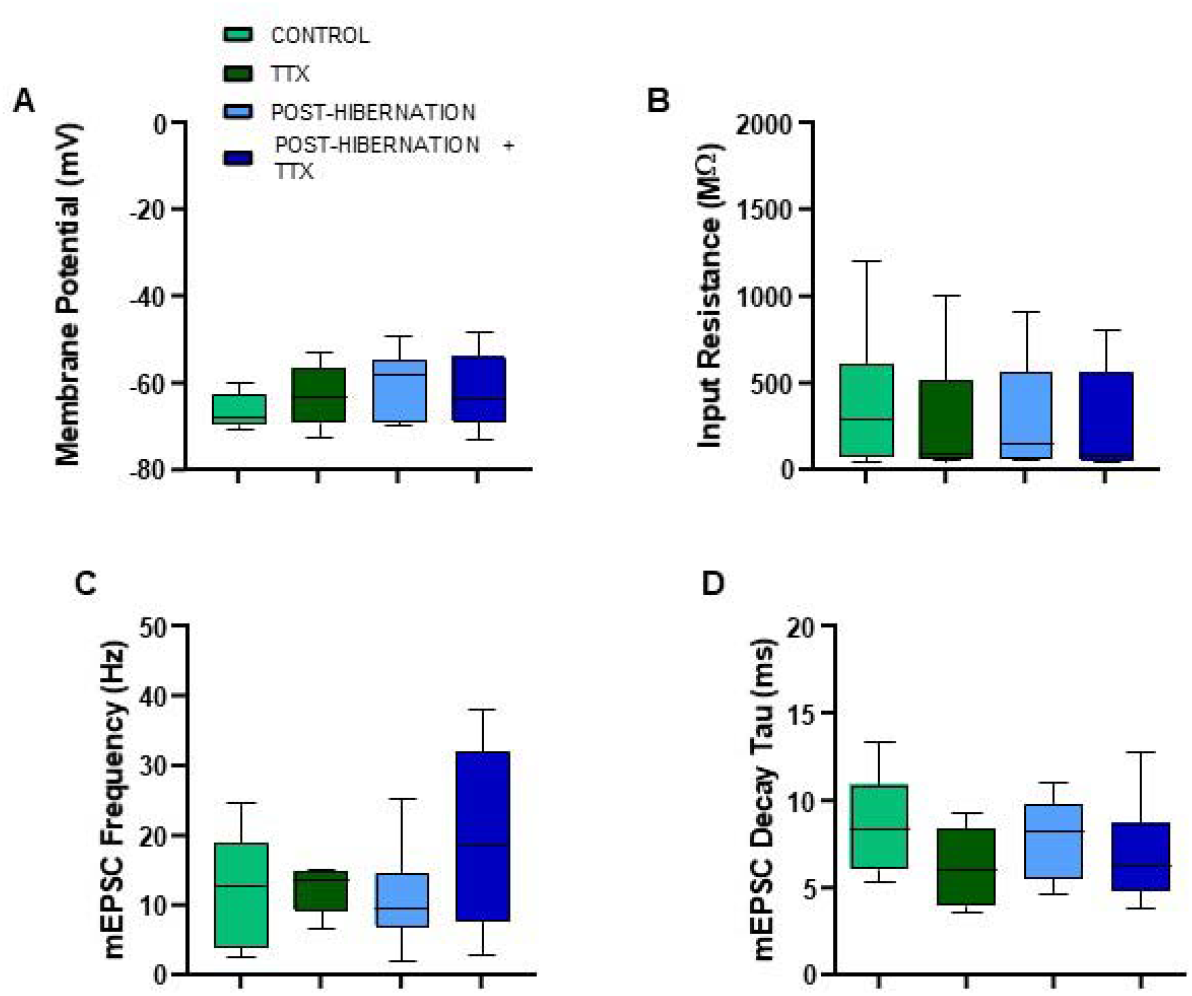
Membrane and other mEPSC properties are not affected following hibernation with or without activity in the post-hibernation period. A. Membrane potential, p=0.081 (one-way ANOVA); B. Input resistance, p=0.676 (Kruskal-Wallis test); C. mEPSC frequency, p=0.403 (Kruskal-Wallis test); D. mEPSC decay time constant, p=0.061 (Kruskal-Wallis test). Control (n=18), TTX (n=19), post-hibernation (n=17), post-hibernation+TTX (n=16). Box and whisker plots show the interquartile range. Whiskers extend to maximum and minimum values.

To upregulate synaptic strength in an inactivity-dependent manner, we hypothesized that reduced Ca^2+^ influx through voltage-gated Ca^2+^ channels may play a role. Thus, we exposed rhythmic circuit preparations to 10 μM nimodipine, a blocker of L-type Ca^2+^ channels in motoneurons of this system^30^ for 16 hrs. Exposure of warm-acclimated preparations to nimodipine did not affect mean mEPSC amplitude or the distribution relative to warm-acclimated time controls (Fig. 5A-B). Therefore, block of L-type Ca^2+^ does not mimic the synaptic response to TTX and hibernation. Posthibernation preparations exposed to nimodipine still resulted in upregulation of cell-averaged mEPSCs compared to time controls (Fig 5A-B), just as we observed with TTX, hibernation, and hibernation+TTX. When comparing the post-hibernation+nimodipine group to post-hibernation alone, cell-averaged mEPSC amplitude were not different (Fig. 5C). Overall, reduced L-type channel function does not drive compensation in this system.

**Figure 5.**
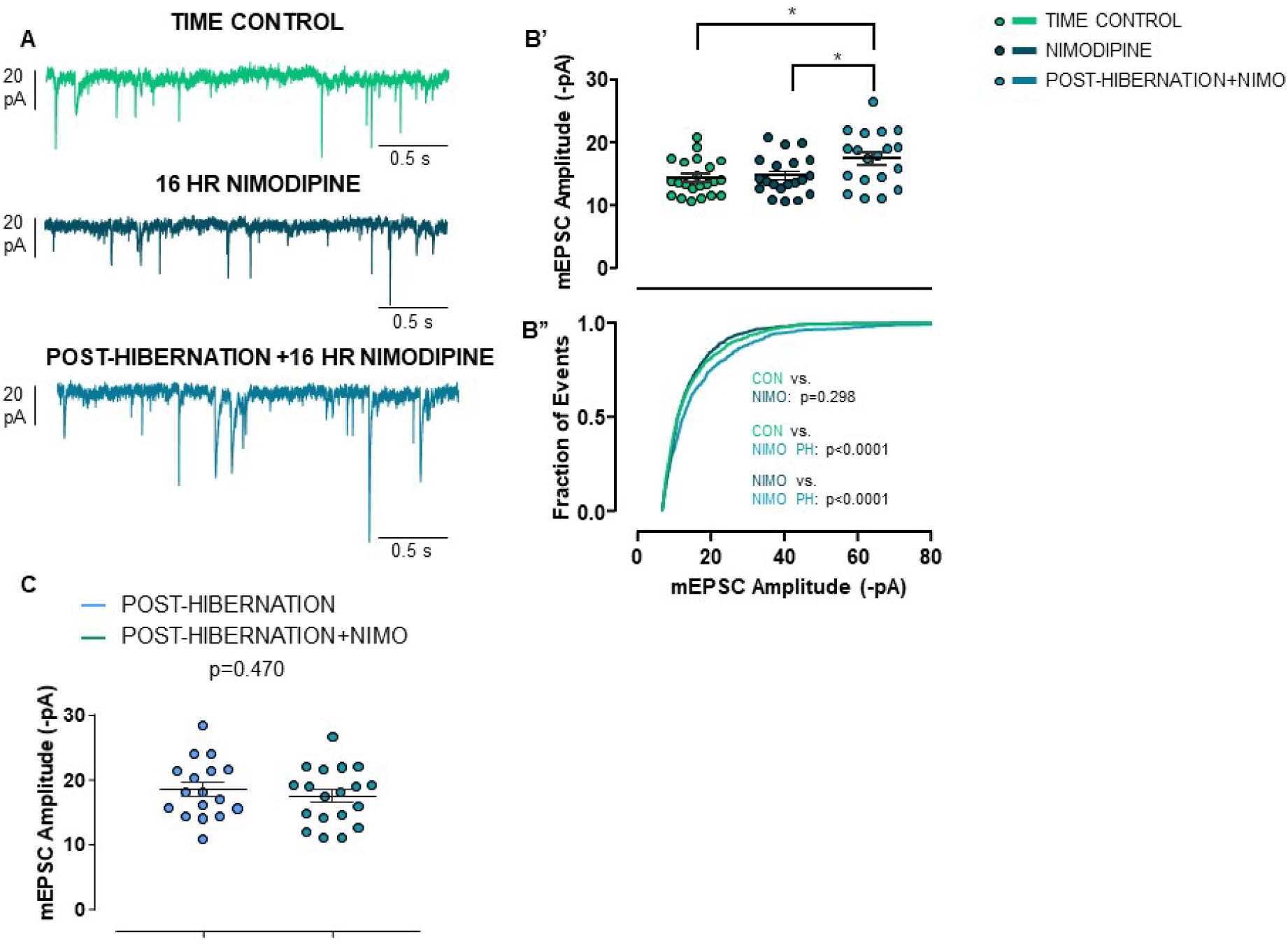
Block of L-type Ca^2+^ channels does not increase mEPSC amplitude. (A) Example mEPSC traces recorded in whole-cell voltage clamp in time control (n=22 from 6 animals), 10 μM nimodipine (n=20 from 5 animals), and post-hibernation+nimodipine (n=19 from 5 animals). (B’) Cell-averaged mEPSC amplitudes and (B”) cumulative distribution of mEPSC amplitudes. Nimodipine exposure does not increase mean synaptic strength or alter the distribution of mEPSC amplitudes. After hibernation, exposure to nimodipine still results in enhanced synaptic strength. (C) Post-hibernation (n=17 from 5 animals) and post-hibernation+ nimodipine (n=19 from 5 animals) have similar group means (unpaired t test).

Given that the entire mEPSC amplitude distribution is thought to be regulated by a multiplicative scaling factor during inactivity^24^, we tested whether the increase in postsynaptic strength in each of our experimental conditions increases according to a multiplicative scaling rule. We used the rank-order method to test for synaptic scaling of mESPC amplitudes in populations of neurons^4,8,31^. To do this, mEPSCs from control and treatment groups (TTX, hibernation, hibernation+TTX, hibernation+nimodipine) were rank-ordered from the smallest to largest amplitude, plotted against each other (ranked control vs. ranked treatment), and fit with a linear regression (Supplemental Fig. 1). The treatment distribution for each form of inactivity was then mathematically downscaled by the equation of the line, and the scaled distribution was compared to control. Evidence for multiplicative scaling is usually taken as a statistically indistinguishable matching between the scaled and control distributions of mEPSC amplitudes^4,12,32^.

All treatments strongly right-shifted the amplitude distribution of mEPSCs relative to controls, as expected based on the mean upregulated synaptic strength for all conditions (Fig. 6A-E; K-S test; time control vs. all treatment conditions; p<0.0001). When we performed the linear transformation to downscale each treatment distribution, interestingly, not all showed evidence of multiplicative synaptic scaling. Hibernation and hibernation+TTX, led to multiplicative scaling (Fig. 6B-C), but TTX alone (Fig. 6A) and hibernation in the presence of nimodipine (Fig. 6D-E) did not. These results suggest that hibernation upregulates synaptic drive while maintaining the balance of synaptic weights according to a scaling rule as we previously showed^15^ The multiplicative organization of compensation appears to be maintained by Ca^2+^ signaling through L-type channels, independent from factors that increase mean synaptic strength, as blocking voltage-gated Ca^2+^ channels disorganized multiplicative scaling but did not alter the mean increase in synaptic drive.

**Figure 6.**
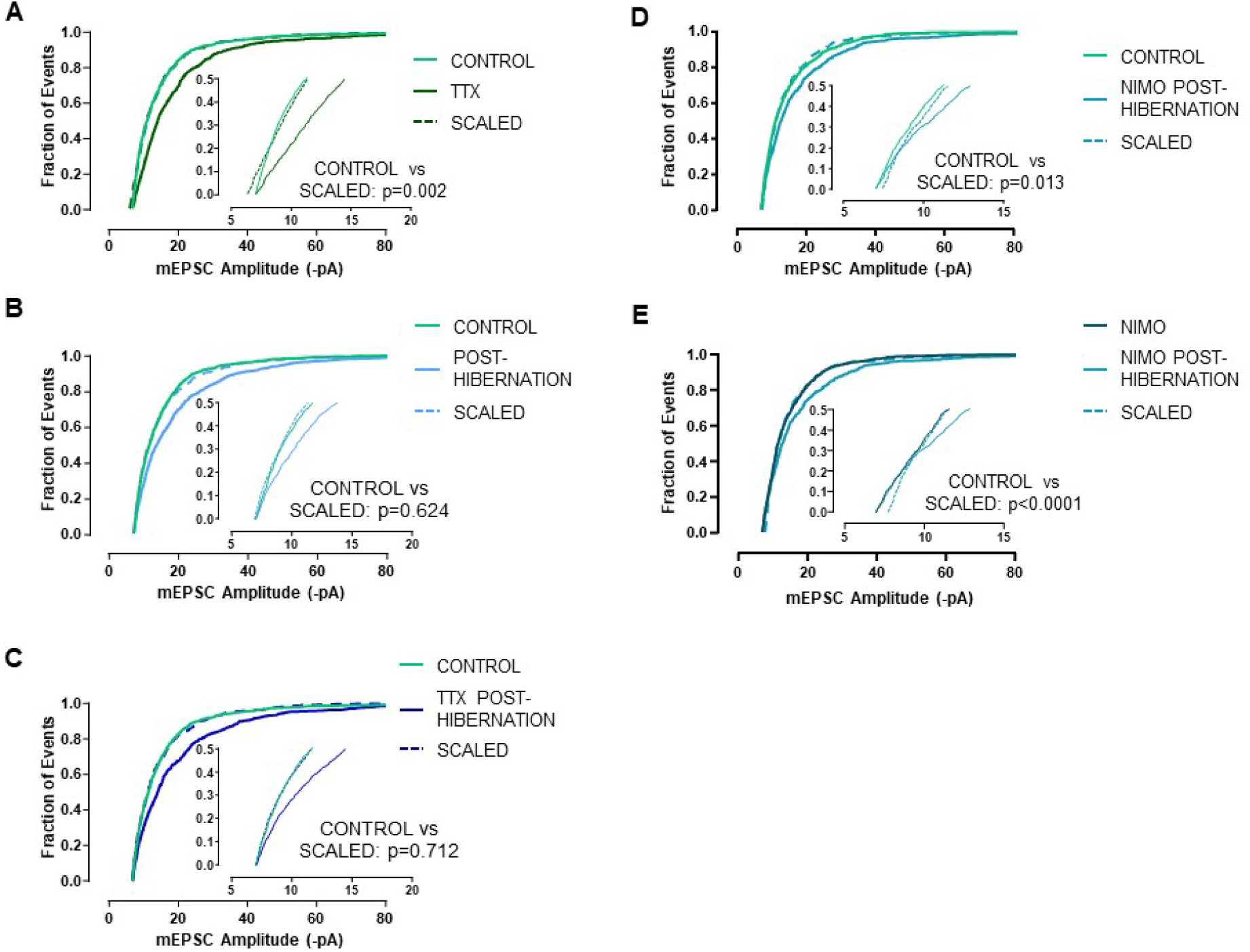
Hibernation, but not TTX, upregulate synaptic strength through multiplicative scaling, and multiplicative scaling is maintained by Ca^2+^ feedback. Cumulative distributions of mEPSC amplitudes for time control, treatment (TTX, posthibernation, post-hibernation+TTX, post-hibernation+nimodipine), and mathematically downscaled treatment distributions. TTX does not lead to precise scaling of mEPSC amplitudes (A). Upregulated TTX mEPSCs were downscaled by the linear equation y=1.47x-2.21 (B) Hibernation and hibernation+TTX increases mEPSCs through multiplicative scaling (C). Upregulated hibernation mEPSCs were downscaled by the linear equation y=1.53x-3.58 and upregulated mEPSC after hibernation+TTX were downscaled by the equation y=1.61x-4.14. (D-E) After hibernation in the presence of nimodipine, mEPSC amplitudes were no longer increased according to multiplicative scaling. (D) uses time control as the control group and (E) uses the warm-acclimated nimodipine as the control. When using the time control as control, upregulated mEPSC amplitudes after hibernation+nimodipine were downscaled according to the equation, y=1.45x-3.76. When using nimodipine from warm acclimated animals as the control, upregulated mEPSCs were downscaled according to the equation y=1.52-4.75. All comparisons between control and treatment yielded p values <0.0001 (Kolmogorov-Smirnov Test.). P values shown in the figures compare control vs. scaled distributions (Kolmogorov-Smirnov Test).

It has been recently suggested that plotting the ratio of treatment to control across the distribution of mEPSCs provides another way to assess if scaling is uniformly multiplicative^40^ In this case, a flat line across the entire distribution would suggest scaling is multiplicative. Therefore, we plotted the ratio of treatment/control vs. control for each experimental group (Supplemental Fig. 2). Consistent with the linear transformation method, both hibernation data sets showed a roughly flat ratio line relative to situations where scaling has been deemed “non-uniform”^33,34^ The pattern for both hibernation groups differ visually from TTX alone (green) and post-hibernation+nimodipine (light green), with post-hibernation+nimodipine deviating substantially from uniform scaling (Supplemental Fig. 2). These results suggest hibernation consistently increases the mEPSC distribution in a roughly multiplicative manner and Ca^2+^ signaling maintains multiplicative organization after hibernation.

To infer the role of these synaptic modifications on network function, we assessed circuit activity of the intact respiratory network. The respiratory network is a central pattern generating network which permits the recording of these motor neurons as they receive their normal presynaptic inputs *ex vivo* (Fig. 7A). In this preparation, extracellular population motor burst frequency of the cut vagus root provides information regarding the function of the rhythm generating networks^35^, and integrated burst shape relates to motoneuron firing and/or recruitment that drives contraction of respiratory muscles^36^ Thus, we could observe physiological output from motoneurons under conditions where synaptic strength did not change (control and warm-nimodipine), where it was upregulated and multiplicatively scaled (post-hibernation), and upregulated but not correctly scaled (post-hibernation+nimodipine). Across controls, post-hibernation, and post-hibernation+nimodipine, no differences existed in frequency of the respiratory rhythm at 13 hrs *in vitro* (one-way ANOVA, p=0.3215). However, preparations treated with nimodipine for 13 hrs after hibernation had shorter motor burst widths relative to all other groups (Fig. 7A-B). Burst width narrowing appeared to be a chronic effect induced by nimodipine following hibernation, as acute application did not affect burst width before and after hibernation, and burst width without nimodipine (control or post-hibernation) did not change with time (Fig. 7C). In addition, membrane potential, input resistance, and action potential firing frequency of laryngeal motoneurons were not different across groups, indicating chronic nimodipine does not induce changes in intrinsic membrane properties (Fig. 7D-F). Overall, these experiments suggest that inactivity increases synaptic strength in response to hibernation and proper scaling of the mESPC distribution appears to be important for normal network function associated with breathing after hibernation.

**Figure 7.**
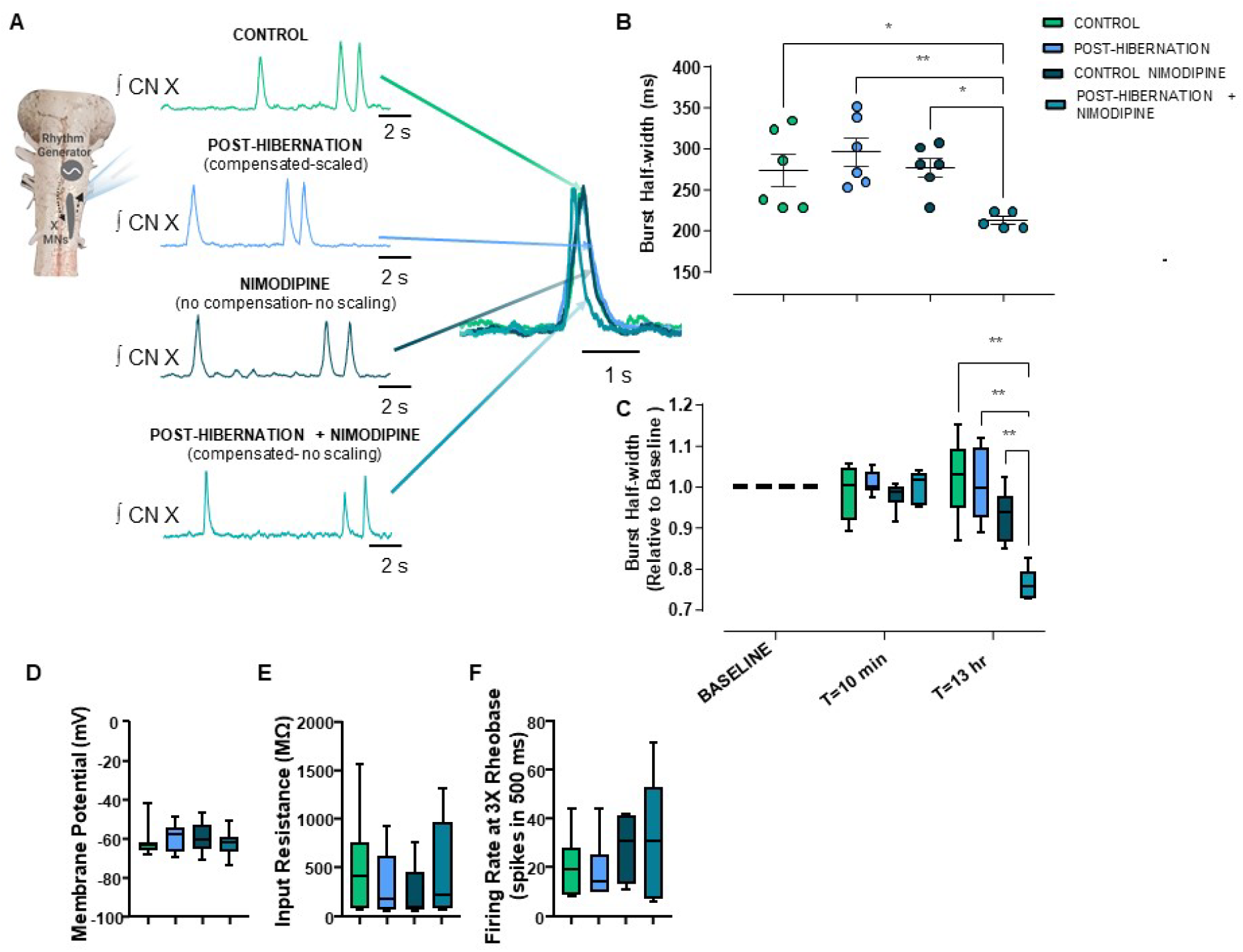
A failure to precisely scale motoneuron synaptic strength is associated with reduced respiratory motor outflow. (A) Simplified schematic of the brainstem preparation, highlighting integrated respiratory-related motor output recorded from the vagus nerve. The four groups shown are time control, post-hibernation (associated with scaled compensation), nimodipine in warm-acclimated control (no compensation, no scaling), and post-hibernation+nimodipine (unscaled compensation) for 13 hrs *in vitro.* Traces are compressed or expanded and overlayed to highlight differences across groups. (B) Nimodipine applied following hibernation resulted in narrower respiratory bursts from the vagus nerve (One-way ANOVA with Holm-Sidak’s Multiple Comparisons Test; control vs. post-hibernation nimo, p=0.049, post-hibernation vs. posthibernation nimo, p=0.007, nimo vs posthibernation nimo=0.041). (C) Nimodipine resulted in a slow, reduction of burst width in the post-hibernation group, as acute exposure to nimodipine did not affect burst amplitude in any group (Two-way ANOVA followed by Holm-Sidak’s multiple comparisons test). However, burst width decreases with longer exposure of nimodipine only after hibernation. n= 6,6,6, and 5 brainstem preparations for control, post-hibernation, nimodipine, and post-hibernation nimodipine, respectively. Intrinsic membrane properties were not different across groups. Membrane potential (D, p=0.356; Kruskal-Wallis test), input resistance (E, p=0.409; Kruskal-Wallis test), firing frequency at 3X rheobase (F; p=0.504; one-way ANOVA). For (A), n= 6,6,6, and 5 brainstem preparations for control, post-hibernation, nimodipine, and post-hibernation nimodipine, respectively. For (D&E), n= 20, 17, 20, and 19 neurons for control, posthibernation, nimodipine, and post-hibernation+nimodpine, respectively. For assessment of firing rate (F), n= 7,6,6,7 neurons were used in control, post-hibernation, nimodipine, and post-hibernation+nimodpine, respectively. Box and whisker plots show the interquartile range. Whiskers extend to maximum and minimum values *p<0.05, **p<0.01.

## Discussion

Neural circuits can compensate for activity perturbations with the apparent goal of regulating neuronal output. American bullfrogs must produce appropriate motor drive for breathing after potentially months of network inactivity during hibernation. Our previous results showed that scaling up the strength of motoneuron synapses helps promote motor output after hibernation to generate breathing. Data here support the idea that inactivity during hibernation drives up synaptic strength and that Ca^2+^ signaling independently controls the precise distribution of upregulated synaptic weights. Overall, these results suggest that multiple signals interact to regulate synaptic compensation in response to environmental challenges *in vivo.*

We previously showed that synaptic strength is increased at ~2 months of hibernation^15^. Here we demonstrate that increases in synaptic drive are present at 2 weeks into aquatic hibernation, which saturate a new steady-state level (Fig. 1C). These responses were comparable to short-term inactivity by TTX (Fig. 2C) and are therefore consistent with upregulation of synaptic strength in response to a signal related to inactivity. Supporting this conclusion, TTX did not cause further increases in synaptic strength when applied after hibernation, suggesting the mechanisms involved in enhancing synaptic strength were already engaged by hibernation. In this regard, inactivity may upregulate synaptic strength through mechanisms that involve the loss of spiking, reduced AMPA receptor activation, and reduced neuromodulation^21,37,38^ That said, synaptic compensation does not seem to strictly follow negative feedback regulation, as inactivity over vastly different time scales (16 hours to 1 month) produced similar degrees of mean synaptic strengthening. Additionally, synaptic strength remained elevated despite the return of activity for at least 16 hrs after hibernation and did not differ when activity was present or absent during the post-hibernation period. Thus, our results indicate that inactivity likely drives up synaptic strength, but a more complicated regulation scheme appears to influence the maintenance of this plasticity.

L-type Ca^2+^ channels are attractive mediators of synaptic compensation, as they communicate changes in intracellular Ca^2+^ to the nucleus and other intracellular signaling pathways during inactivity^22,25^ In contrast, chronic block of L-type Ca^2+^ channels did not affect the mean mEPSC amplitude. Although our data rule out the possibly for a role of nimodipine-sensitive L-type Ca^2+^ channels in compensation, other Ca^2+^ channels could contribute. For example, in respiratory motor neurons of this species, nimodipine blocks the 30-40% of the high-voltage activated Ca^2+^ current^30^, indicating the expression of other somatodendritic calcium channels on respiratory motoneurons. Nevertheless, L-type Ca^2+^ channels do not appear to drive synaptic compensation, differing from traditional models of homeostatic plasticity in neuronal cultures^22,25,26,39^.

Synaptic inputs are often upregulated across the entire distribution by the same multiplicative factor^4^. An interesting finding of this study is hibernation, but not TTX, exhibited multiplicative scaling when using a linear transformation of ranked mEPSCs^4,8,14^ While we present evidence for and against multiplicative scaling under different conditions, we note recent studies suggesting complexity in this interpretation. Synaptic spine density and AMPAR expression at individual synapses do not scale uniformly within the same neuron even though multiplicative scaling appears to occur at the population level^40,41^. A recent study also suggested that TTX may not lead to multiplicative scaling in different mammalian neuronal types when assessed using the linear transformation method^34^, although significant evidence points to the contrary^12,22,42,43^ Consistent with multiplicative scaling as assessed by the linear transformation (Fig. 6), the ratio of the post-hibernation:control amplitude plotted as a function of control amplitude is roughly flat relative to other systems where scaling has been considered “non-uniform”^33,34^ (Supplemental Fig. 2). Although both lines of evidence point to multiplicative scaling, we acknowledge that the ratio line is not perfectly straight. This likely occurs in part because the “rank-ratio method”^34^ does not account for values on the bottom of the treatment distribution that lack a partner in the control distribution due to the mEPSC detection cutoff. Misalignment due to the cutoff, therefore, can make perfectly uniform scaling appear non-uniform^34^ The linear transformation method controls in part for this issue through the y-intercept term^4,31,32^ so long as the mEPSCs are evenly spread across various amplitudes,^41^ which largely holds for our data (Supplemental Fig. 1). Collectively, these results support the idea that hibernation drives up synaptic strength while maintaining the balance of synaptic weights, while inactivity without hibernation upregulates synaptic strength without multiplicative scaling.

Our results suggest that a separate factor associated with hibernation controls the multiplicative nature of synaptic compensation, as inactivity without hibernation does not follow the same pattern (Fig. 6). We find that blocking L-type Ca^2+^ channels *following* hibernation disrupted multiplicative scaling without influencing the cell-average amount of compensation (Fig. 6). These results suggest that Ca^2+^ feedback maintains multiplicative scaling after hibernation, and these processes are separate from those that upregulate mean synaptic strength in response to inactivity in the first place. Given that we observed evidence for multiplicative scaling after hibernation in the presence of TTX (Fig. 6), the sum of our data suggests maintenance of multiplicatively scaled mEPSC distribution is Ca^2+^-dependent but not activity-dependent. As intracellular calcium transients tend to follow patterned activity, these results may appear contradictory. However, spontaneous neurotransmitter release in silent neurons can activate L-type channels at subthreshold voltages to induce neuronal plasticity^44,45^ As cold temperature increases membrane input resistance^46,47^, spontaneous neurotransmitter release may cause larger postsynaptic depolarizations compared to warm animals, generating significant Ca^2+^ influx through L-type channels to orchestrate scaling during the winter. This interpretation diverges significantly from classic models of synaptic scaling through a multiplicative factor that drives up synaptic strength during inactivity. Instead, our data suggest that inactivity during winter upregulates synaptic strength (Fig. 2–3) through mechanisms that do not involve multiplicative scaling (Fig. 6A), and the winter state engages a separate regulatory system involving L-type Ca^2+^ channels to “rebalance” the mEPSC distribution (Fig. 6D-E). Although we did not observe differences in rhythmic burst frequency of the network in nimodipine, it is possible that chronic nimodipine altered the firing pattern of motoneurons during ongoing rhythmic activity, and subsequently modified the scaling arrangement after hibernation. These parallel pathways warrant further investigation and may be relevant to complex physiological perturbations in other systems.

Our previous study identified that upregulation of synaptic strength acts to maintain motor output associated with breathing following hibernation^15^ Here we present evidence that proper organization of the mEPSC amplitude distribution influences network function, as failure to correctly scale synaptic inputs after hibernation reduced respiratory motor output (Fig. 7). We recognize the challenge of attributing changes in network-level properties to shifts in the distribution of synaptic weights, and we made several attempts to address other variables that could explain these results. First, rhythmic burst frequency of the network was the same across all groups *ex vivo;* thus, overall activity differences are unlikely to explain shorter motor bursts. Second, several intrinsic membrane properties were not different across scaled and “non-scaled” conditions (Fig. 7D-F), suggesting that changes in excitability did not weaken the motor outflow. Third, warm-acclimated control preparations exposed to nimodipine did not affect any other electrophysiological property, arguing against actions of nimodipine on different parts of the network (Fig. 6–7). Finally, acute exposure to nimodipine did not affect burst duration in any group (Fig. 7C), suggesting that chronic nimodipine exposure after winter reduces motor outflow by disrupting the mEPSC amplitude distribution. All of this said, we cannot state with certainty if narrower respiratory bursts reflect improperly organized synapses on motoneurons (Fig. 6D-E) or actions of nimodipine on premotor neurons that regulate motor outflow in a hibernation-dependent manner. Nevertheless, these results, in concert with our previous findings^15^, indicate that synaptic scaling is important for regulating motor performance shortly after inactivity in hibernation.

In conclusion, synaptic scaling has clear adaptive value following hibernation in the frog, as it ensures adequate motor drive for lung ventilation^15^ (present study Fig. 7). More broadly, our results suggest that homeostatic synaptic plasticity could involve a more complicated regulation scheme when implemented in living animals during natural perturbations. Synaptic compensation is often viewed as a conceptually simple feedback process, whereby neurons sense their activity state and elicit appropriate compensation in attempt to regulate network output. In contrast, our data suggest that different environmental variables may control different aspects of the same compensation response. Many neurological disorders are thought to involve a failure of homeostatic synaptic and intrinsic mechanisms^9^, all of which occur in complex organisms with diverse types of environmental feedback onto the nervous system. Our results emphasize that confronting how the environment shapes compensation will be necessary to understand the expression and control of homeostatic compensation. Learning from animals that use these mechanisms to survive extreme environments, therefore, has the potential to provide new insight into the regulation of neuronal function in both health and disease states.

## Materials and Methods

### Ethical Approval and Compliance

All animal treatments and protocols were approved by the Institutional Animal Care and Use Committee (IACUC) at the University of North Carolina at Greensboro (protocol #19-006). All methods reported are in accordance with the IACUC at University of North Carolina-Greensboro, as well as the ARRIVE guidelines.

### Animals

Bullfrogs of either sex weighing approximately 100 g were purchased from Rana Ranch (Twin Falls, ID, USA). Control frogs were maintained at 20 °C in plastic tubs and were provided aerated water with access to wet and dry areas. Frogs were fed pellets provided by Rana Ranch once per week. Hibernation conditions were simulated in temperature controlled incubators. The temperature of water was lowered from 20°C to 2°C over the course of a week, dropping about 2-3 degrees per day. On the seventh day, a plastic screen was placed at the top of the water to prevent access to the surface. Animals do not eat at cold temperatures; therefore, food was also withheld while frogs remained at 2°C. Frogs in each condition were kept on a 12 hr:12 hr light: dark cycle. For all experiments, frogs were randomly assigned to each experimental group upon delivery. Experiments were only performed during the light cycle.

### Dissection and Tissue Preparation

Brainstem-spinal cord dissections were performed as previously described^15^ Briefly, frogs were deeply anesthetized with approximately 1 ml of isoflurane per liter and rapidly decapitated. The head was immersed in chilled bullfrog artificial cerebrospinal fluid (aCSF) and the brainstem-spinal cord was removed, keeping the motor rootlets of the vagus nerves intact (aCSF; concentrations in [mM]: 104 NaCl, 4 KCl, 1.4 MgCl_2_, 7.5 glucose, 40 NaHCO_3_, 2.5 CaCl_2_ and 1 NaH_2_PO_4_, and gased with 98.5% O_2_, 1.3%CO_2_; pH = 7.85). Following the dissection, the brainstem was superfused with aCSF with a flow rate of ~6 mL per minute.

To selectively label laryngeal branch motoneurons that innervate the glottal dilator, the 4^th^ branch of the vagus nerve was isolated and then backfilled tetramethylrhodamine dextran 3000 MW (Life Technologies Corporation, Eugene, OR USA). Approximately 1 uL of 10% dextran was loaded into the tip of a glass pipette that fit snuggly around the cut 4th root of the vagus, as this nerve contains axons that mainly innervate the glottal dilator see references provided in^15^. Backfills lasted two hours and produced robust labeling of neurons for identification prior to electrophysiological recording. Following the dye loading process, tissue was sliced with a vibratome at 300 μM. Sections approximately 300-600 μm rostral to the obex were used for electrophysiological recording. Tissue slices were left to recover for an hour before electrophysiological recording.

### Whole-cell patch clamp recordings

Slices containing labeled motoneurons were transferred to the recording chamber, stabilized with a nylon grid, and exposed to aCSF at a flow rate of ~1-2 mL per min supplied via a gravity-fed system. The region containing labeled neurons was imaged at 4x, and then individual neurons were located within the slice at 60x (excitation, 540 nm: emission, 605 nm). Imaging was performed with a Hamamatsu ORCA Flash 4.0LT sCMOS Camera (Hamamatsu Photonics, Japan). Identified neurons were approached with the whole-cell patch pipette controlled by a Sutter Instruments micromanipulator system, which was equipped with a MP-285 micromanipulator and a MPC-200 controller (Sutter Instruments, CA, USA). A P87 horizontal pipette puller (Sutter Instruments, Novato, CA, USA) was used to pull patch pipette, which had resistances of 2-3 MΩ when filled with solution containing the follow composition (in mM): 110 K-gluconate, 2 MgCl2, 10 HEPES, 1 Na2-ATP, 0.1 Na2-GTP, 2.5 EGTA. Pipettes were attached to a headstage connected to an Axopatch 200B amplifier, with signals digitized at 100 kHz using a Digidata 1550 (Molecular Devices, San Jose, California). The cell of interest was approached with gentle positive pressure on the pipette. When an indentation could be seen on the cell in the bright field with IR-DIC optics, positive pressure was removed and a >1 GΩ was formed by applying slight negative pressure through the pipette by mouth. Whole-cell access was obtained by rapid negative pressure applied by mouth to rupture the GΩ seal. For measurements of excitatory postsynaptic currents, cells were voltage clamped at −80 mV to record synaptic currents mediated by AMPA receptors^15^ When spontaneous excitatory synaptic currents (sEPSCs) were recorded in aCSF, no TTX was included during the recording. When miniature excitatory postsynaptic currents (mESPCs) were recorded, 500 nM TTX was added the aCSF. A +10 mV step was periodically performed to estimate series resistance (peak of the transient current) and input resistance (steadystate current) using Ohm’s law. Measurements of membrane potential were made in current clamp, and firing rate was assessed at 3X rheobase in some current clamp experiments before application of TTX to measure mEPSCs. We often recorded more than one neuron per slice; however, TTX does not have a fast washout time in this preparation; therefore, firing properties in current clamp were not assessed for every cell. Previous work in these neurons suggests that firing response to current injection does not change following hibernation^15^, and our results in this study, despite a smaller sample size, support this previous finding.

### Extracellular recordings

Extracellular output from the cut vagus nerve was recorded with a bipolar suction electrode connected to an A-M Systems Model 1700 amplifier (bandpass filtered between 10-1000 Hz, 100X gain). Activity of this nerve is primarily generated by the 4^th^ branch of the vagus nerve (the same neurons we studied in slices), and provides the main drive to the glottal dilator, which gates airflow into and out of the lung in anuran amphibians^48^

### Experimental Treatments

Three main series of experiments were performed in this study: (1) those that varied the duration of cold acclimation and then assessed synaptic function immediately after removing the animal from the hibernation environment, (2) those that used either warm-acclimated control animals or 1 month hibernation animals and then exposed these brainstem preparations to TTX for 16 hrs in vitro, and (3) those that used either warm-acclimated control animals or 1 month hibernation animals and exposed each to 10 μM nimodipine for 16 hrs. For series 1 experiments control frogs (20°C) or cold-acclimated (2°C for 2 or 4 weeks) were used, and then synaptic function was assessed after animals were removed from these environments to roughly determine the time course of synaptic compensation. In series two experiments our main goal was to compare to post-hibernation synaptic function to inactivity with 100 nM TTX. In addition, we assessed synaptic function in post-hibernation animals after silencing rhythmic motor output with TTX. TTX silenced all preparations for the entire 16 hr period (data not shown). In series three experiments, we exposed warm-acclimated control and posthibernation rhythmic preparations to 10 μM nimodipine for 16 hrs to block L-type Ca^2+^ channels, a dose that blocks these channels in frog respiratory motoneurons^30^, and a separate time control group was run in parallel. Following these treatments all mEPSC recordings were performed in standard aCSF with 500 nM TTX.

### Data Analysis

All data analysis was performed using the peak analysis module on LabChart 8 (ADInstruments, Sydney, Australia). sEPSC and mESPCs events were recorded for 30 seconds and averaged to estimate excitatory synaptic drive due to AMPAR transmission. Cutoff amplitude was set at 7 pA. All analyses were checked by eye to verify EPSCs were correctly identified and were not performed blinded to treatment. To infer synaptic scaling, we used the standard rank-order method as originally described by Turrigiano et al.^4^ Briefly, a series of 50 mESPCs were randomly chosen for each cell in the group and the entire distribution was ranked from lowest to highest for control and treatment conditions. To assess synaptic scaling, the same number of synaptic events must be present in the control and treatment group. However, in some cases in the present study, control and treatment conditions had different numbers of cells (typically differing 1-3 neurons). In these situations, cells from groups with the higher number were randomly removed for the scaling analysis by assigning a number to each cell and then using a random number generator chose which cell(s) would be removed. This allowed equal number of mESPCs to be ranked and scaled in each group. Ranked data were plotted and then fit by a linear regression to obtain the slope of the line and intercept for the entire population. ~5-10 individual mEPSC events in each group showed extreme values that deviated from the linear fit. These events may be caused by activation of subsaturated AMPA receptors, as suggested previously^42^, and were therefore removed for the fitting of the ranked data. Each mESPC event in the treatment distribution was then mathematically downscaled according to the the linear equation and plotted as cumulative distributions. We also plotted ranked data as “treatment/control vs. control” across the mEPSC distribution for each experimental group^34^. Input and series resistances were calculated using Ohm’s law by making a 10 mV step, from −80 mV to −70 mV and measuring the resulting transient and stead-state currents, respectively. Series resistance did not vary across groups, and cells with series resistance of >25MΩ were discarded.

### Statistics

Data were analyzed and plotted using GraphPad Prism. Data sets that followed a normal distribution were analyzed using parametric statistical tests. Comparisons between two groups were made using a two-tailed unpaired t test, more than two groups were compared using a one-way ANOVA followed by Holm-Sidak’s multiple comparisons test. When standard deviations were different between groups, a Welch’s test was applied. Some data sets involving three or more comparisons did not follow a normal distribution. In these cases, the Kruskal-Wallis test was used. When comparing two cumulative distributions, the Kolmogorov Smirnov test was run. Brainstem schematics were edited and created with Biorender.com. Significance was accepted at p<0.05.

## Acknowledgements

This research was funded in part by the Department of Defense (W911NF2010275), the National Institutes of Health (1R01NS114514 and 1R15NS112920), and lab startup funds from UNC-Greensboro to J.M.S.

## Author Contributions

JMS conceived and designed the project. TZ performed experiments in Figures 1–4, 6–7. LA performed experiments in Figures 5, 6–7., TZ, LA, and JMS analyzed data. JMS wrote the manuscript. JMS, TZ, and LA edited and approved the final manuscript.

## Data Availability

All datasets generated and/or analyzed during the current study are available from the corresponding author upon reasonable request.

